# Evidence of sensory error threshold in triggering locomotor adaptations in humans

**DOI:** 10.1101/2024.02.19.581021

**Authors:** E. Herrick, S. Yakovenko

## Abstract

Changing body biomechanics or external conditions trigger neural adaptations to optimize motor behavior. While the adaptations appear continuous to minimize movement errors, not all errors necessarily initiate sensorimotor adaptations. The locomotor system exhibits resilience to change and rehabilitation. This suggests the presence of an error threshold to trigger the adaptation mechanism. Here, we imposed kinematic and kinetic constraints of stepping using a passive orthosis and real-time feedback about limb loading. We aimed to manipulate stepping asymmetry and explore the interplay between adaptation and locomotor error magnitude. Uninjured healthy adults were tested in three locomotor conditions: unconstrained, constrained, and washout unconstrained walking. Surprisingly, kinematic asymmetry alone did not induce persistent adaptation. However, the addition of asymmetric interlimb loading triggered the expected adaptation. Our finding suggests that uninjured locomotor systems can cope with a specific range of kinematic asymmetries without initiating persistent adaptations that lead to aftereffects, and that loading may be a key variable for evoking the adaptation. The presence of an error threshold may mitigate possible disruption of vital motor functions and contribute to locomotor adaptation during walking. These insights elucidate the mechanism of neural plasticity and have implications for rehabilitation.

## 2 INTRODUCTION

Human walking is a versatile behavior that adapts not only to unpredictable environments, but also to internal errors within neural and biomechanical mechanisms. Multiple mechanisms participate in reducing errors in walking, a fundamental motor behavior of humans and other terrestrial vertebrates. The difference between planned and executed movement and posture is minimized by viscoelastic muscle responses also known as pre-flexes, sensory feedback actions, and predictive feedforward drive [1,2]. The feedback mechanisms can rapidly compensate for minor and unpredictable external perturbations; yet, the neural pathways involved in pattern generation and coordination may be required to change when body configuration changes, for example, due to aging or trauma. Repeated exposure to a novel mechanical constraint that causes error feedback can engage modifications and gradually optimize movement to new motor demands. The removal of constraint often reveals aftereffects and requires de-adaptation to return the system to the previous optimal state, as illustrated in the Helmholtz’s prism adaptation task [3–6]. Similar to other motor pathways, the locomotor system has been shown to exhibit adaptation and de-adaptation processes, for example, in walking on a split-belt treadmill with asymmetric stepping [7–10]. The study of locomotor adaptations not only provides insight into motor learning [11], but also guides rehabilitation of symmetric walking [12,13].

The prominent computational frameworks explaining the closed-loop movement control is based on the internal models of body dynamics [14]. The inverse models of body dynamics, see (P*)^-1^ element in Figure 1, can transform kinematic signals into motor commands (*efference*). To overcome processing and transmission delays, the collateral feedforward motor signals (*efference copy*) is processed through the internal forward model (P*) to predict and compare sensory feedback during movement execution, as shown in Figure 1. The *external* errors can then be used within the inverse models to generate the automatic compensation to the ongoing motor command. The mismatch (ε) of expected and actual sensory signals identifies unexpected *external* or *internal* errors. The *internal* error may require adaptation of the execution and prediction pathways to reduce the discrepancy following a movement. In the process of motor adaptation, sensory errors drive the process resolving the sensorimotor predictions through the internal representation in the cerebellum, as was shown in reaching tasks [4]. The recalibration of internal model was generally implicated in motor tasks [reviewed in 15] and specifically in the split-belt locomotor studies with large limb speed differences [16,17]. While adaptation of the internal model can optimize body movement in changing conditions reducing overall effort [18,19], there is likely a cost associated with the adaptation and de-adaptation processes. Thus, it is likely that the locomotor system may initiate adaptation only after perceived error exceeds a putative error threshold (ε*) demarcating errors tolerated by the control system and those initiating adaptation within internal representations.

**Figure 1.**
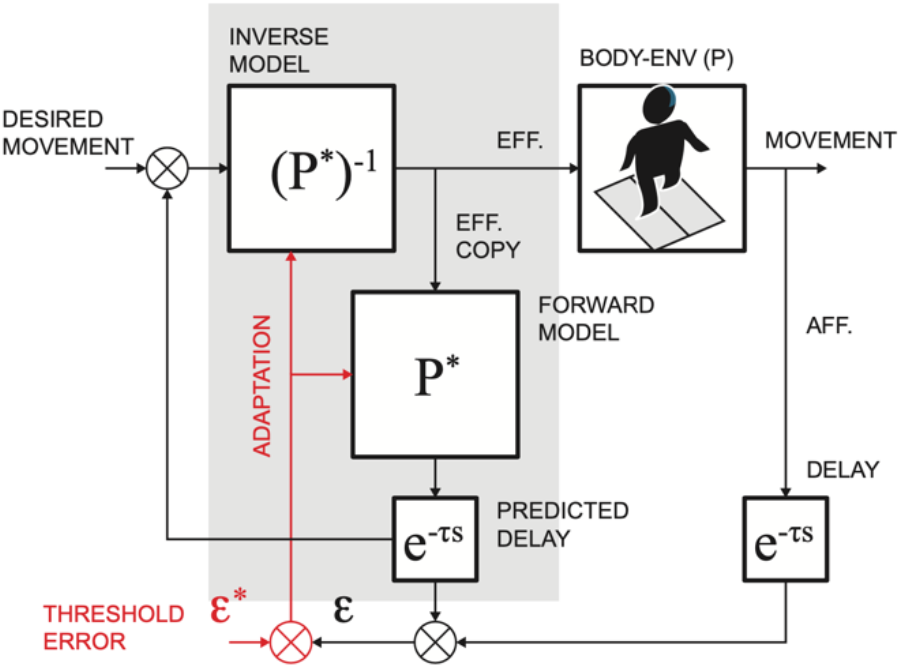
Schematic of the internal dynamic representations for movement execution. P and P* denote respectively body-environment interaction dynamics and its embedded representation. The embedded dynamics in the shaded area can be used for planning and monitoring of movement execution. The hypothetical adaptation pathway (red) may cause changes in dynamic transformations to reduce discrepancy in predicted and observed sensory errors when this difference exceeds a threshold error (ε*).

In this study, we examined the mechanism of locomotor adaptations during asymmetric gait, which has been previously studied to develop effective rehabilitation for individuals with neurological conditions caused by stroke [20,21] and other neurological conditions, for example, parkinsonism [22,23]. Split-belt treadmills were used to impose asymmetrical walking using large speed differences that can trigger the adaptation process and transfer aftereffects to overground walking [16]. However, small limb speed differences are behaviorally related to turning, as shown by experimental studies of walking on a curve [24,25] and simulations [26], and not to the limb preference typically seen in gait pathologies [27]. For this reason, we used kinematic and kinetic constraints to trigger locomotor adaptations. We expected that the asymmetric adaptation would be evident from the persistent aftereffects. We hypothesized that the presence of aftereffects will indicate the recruitment of adaptation mechanism.

## 3 METHODS

### 3.1 Description of population, instrumentation, and experimental procedures

#### Human subjects

23 healthy adults (25.0 ± 6.2 y.o., 12 males, 11 females) with no known neurological disease or persistent musculoskeletal injury consented to participate in the experiment. The participants were divided into two groups: (i) Group 1 (N=11) to perform the walking task with only a kinematic constraint, and (ii) Group 2 (N=12) to perform the walking task with both a kinematic constraint and asymmetric limb loading. The participants walked on the instrumented treadmill with ground reaction force sensors (Bertec, Columbus, OH) while wearing a unilateral stride constraint shown in Figures 2A and 2B. The leg constraint was secured with a strap fastened below the knee protruding anteriorly 2 cm (for Group 1) and 10.5 cm (for Group 2) and medially by 25 cm. This adjustment in protrusion allowed the participant to swing the constrained leg about midway between protraction in our standard constrained condition and the unconstrained condition, decreasing the probability that they would adopt the strategy of walking as if both legs were constrained, see Figure 3B Asymmetric Condition Group 1.

**Figure 2.**
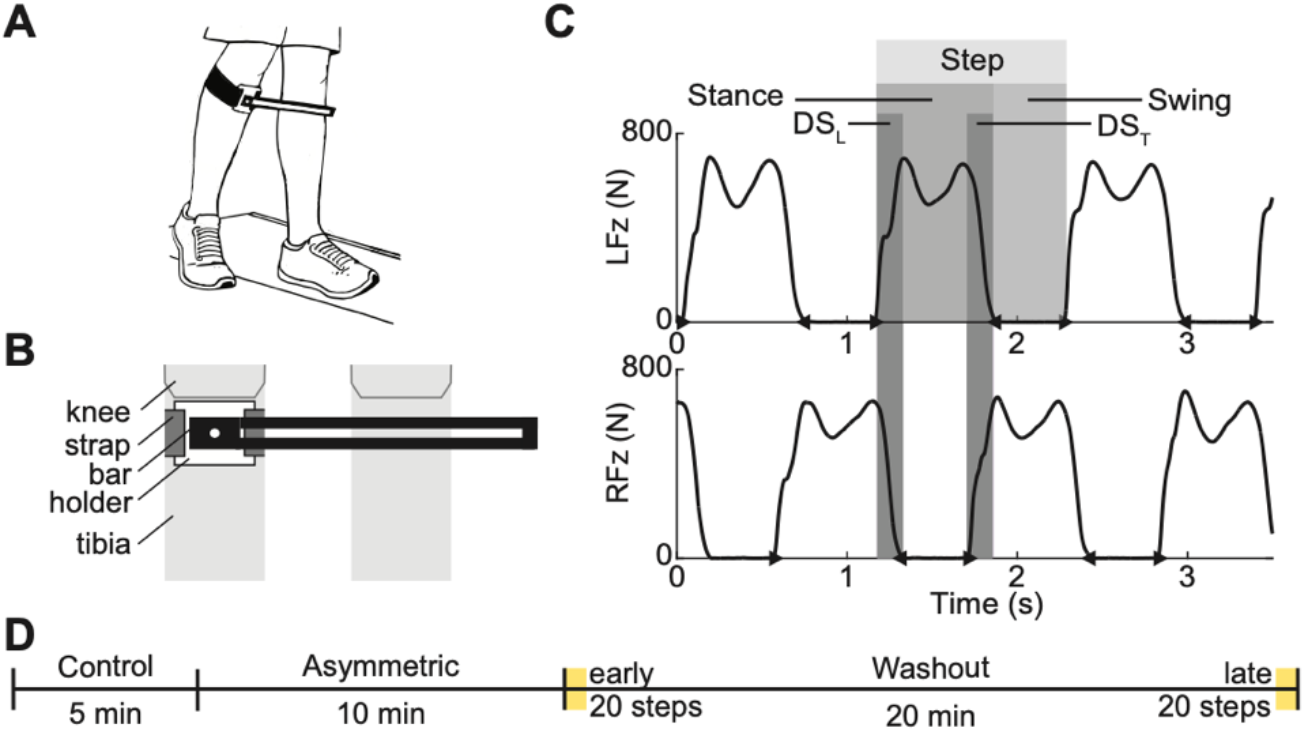
Methodological setup of instrumentation and composition of testing session. **(A)** Depiction of step length constraint. **(B)** Front view of placement of constraint. **(C)** Example of processed ground reaction forces with the detected events defining swing, stance and its subphase termed double stance (DS). **(D)** Session timeline.

**Figure 3.**
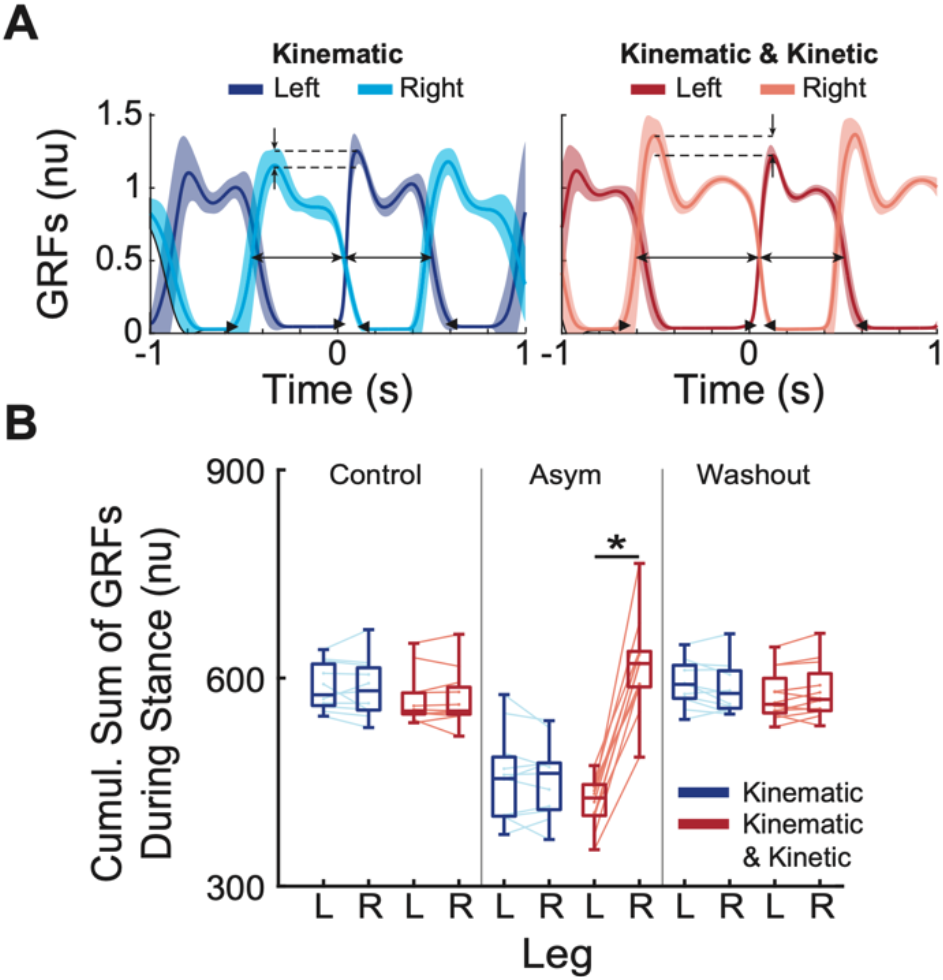
Analysis of the difference in loading symmetry between Groups 1 (*kinematic* constraint only) and 2 (*kinematic* and *kinetic* constraints). **(A)** Examples of vertical GRF profiles for representative participants in each group. Means with the standard deviations of forces were normalized to the participant’s weight and aligned on the onset of stance. **(B)** The cumulative sum of ground reaction forces during stance periods across participants for each leg in every condition. (nu) indicates normalized units. Cumulative sums were normalized to the subject’s weight and the number of steps in each condition. The box plots show the variation across participants for a given condition. The thin lines show the data of individuals within the group with a pair of data points connected by a line representing the difference in loading between the left and right limbs. * Indicates a significant difference with a p *<* 0.05.

Each session consisted of three trial types: control, adaptation (i.e., *Asymmetric*), and washout, see Figure 2D. The control trials documented baseline gait pattern for five minutes of walking at 1 m/s. During the asymmetric trials, participants walked with the device on the right leg constraining the stride length of the left limb. Participants were instructed to not touch the device with the left leg while walking for ten minutes. Specifically for Group 2, participants were explicitly instructed to unload their constrained leg, and verbal feedback was given during the task execution when the unloading was not evident from the real-time vertical component of ground reaction force. The washout period allowed the observation of aftereffects after the removal of the constraints. All procedures were approved by West Virginia University Institutional Review Board.

### 3.2 Data Processing

#### Ground reaction forces and events

Ground reaction forces were sampled at 1000 Hz (National Instruments, Austin, TX). The onsets and offsets of each stance period were detected using a supervised automated thresholding method [28], and the difference between these consecutive events determined stance period. Similarly, swing was defined as the period between the consecutive offset and onset events. The leading and trailing *double stance* sub-phases were defined as the periods when both limbs were on the ground in stance (denoted as *DS*_*L*_, *DS*_*T*_ in Figure 2C, respectively). Because we supervised only a subset of the automatically detected events using an automated threshold detection method [28], an additional standard outlier rejection method was used to remove a small group of unusual events [Tukey’s 1.5 IQR criterion, 29].

#### Asymmetry Index

To quantify the level of asymmetry at each step throughout each condition, we calculated an asymmetry index (AI) based on the double stance subphases for each step. The array of double-stance values was filtered with a moving average (20 samples). The AI was then calculated as the ratio of *leading* and *trailing* double stance periods (*DS*_*L*_, *DS*_*T*_):

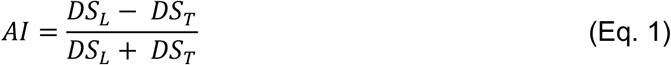

The deviations from zero indicated lateralized gait asymmetry.

#### Aftereffects Analysis

The comparison of asymmetry in the washout period after the removal of constraints was used to test the presence of aftereffects and de-adaptation. We expected that the asymmetry will decrease throughout the washout, which will be evident from comparing AI values of the first and last 20 steps of the washout trial. The Kolmogorov-Smirnov statistic indicated that the difference in AI was non-normally distributed. Consequently, the difference was tested with a one-sided Wilcoxon signed rank test (α = 0.05).

## 4 RESULTS

The asymmetric loading in addition to the kinematic asymmetry was the key variable responsible for the development of aftereffects. Figure 3 shows that participants in Group 2 exhibited asymmetric loading between their left and right limbs (*Asymmetric*: p *<* 0.001) while participants in Group 1 had symmetric loading (*Asymmetric*: 0.6812).

The presence of adaptation in the form of aftereffects was tested with the comparison of asymmetry in early and late periods of the washout condition. Figure 4 shows the comparison of the absolute value of the average AI values across participants in the early (first 20 steps) and late (last 20 steps) portions of the washout condition. For Group 1, the data was not normally distributed (early, p = 0.004, late, p = 0.005). The median for the early washout was smaller than the median for the late washout (0.019 and 0.024) failing a right-tailed Wilcoxon signed rank test (p = 0.29 with α = 0.05). Thus, there was no significant difference between the early and late portions of washout with only a kinematic constraint, see Figure 4A. The average behavior for Group 1 supports this result. In the control condition, participants were nearly symmetrical, with average (± SD) AI values of 0.0216 ± 0.0024. In the asymmetric condition, the imposed asymmetry is evident with average AI values of 0.1706 ± 0.0132. In the washout condition, participants returned to near their baseline with average AI values of 0.0190 ± 0.0038 without any aftereffects from the imposed asymmetry.

**Figure 4.**
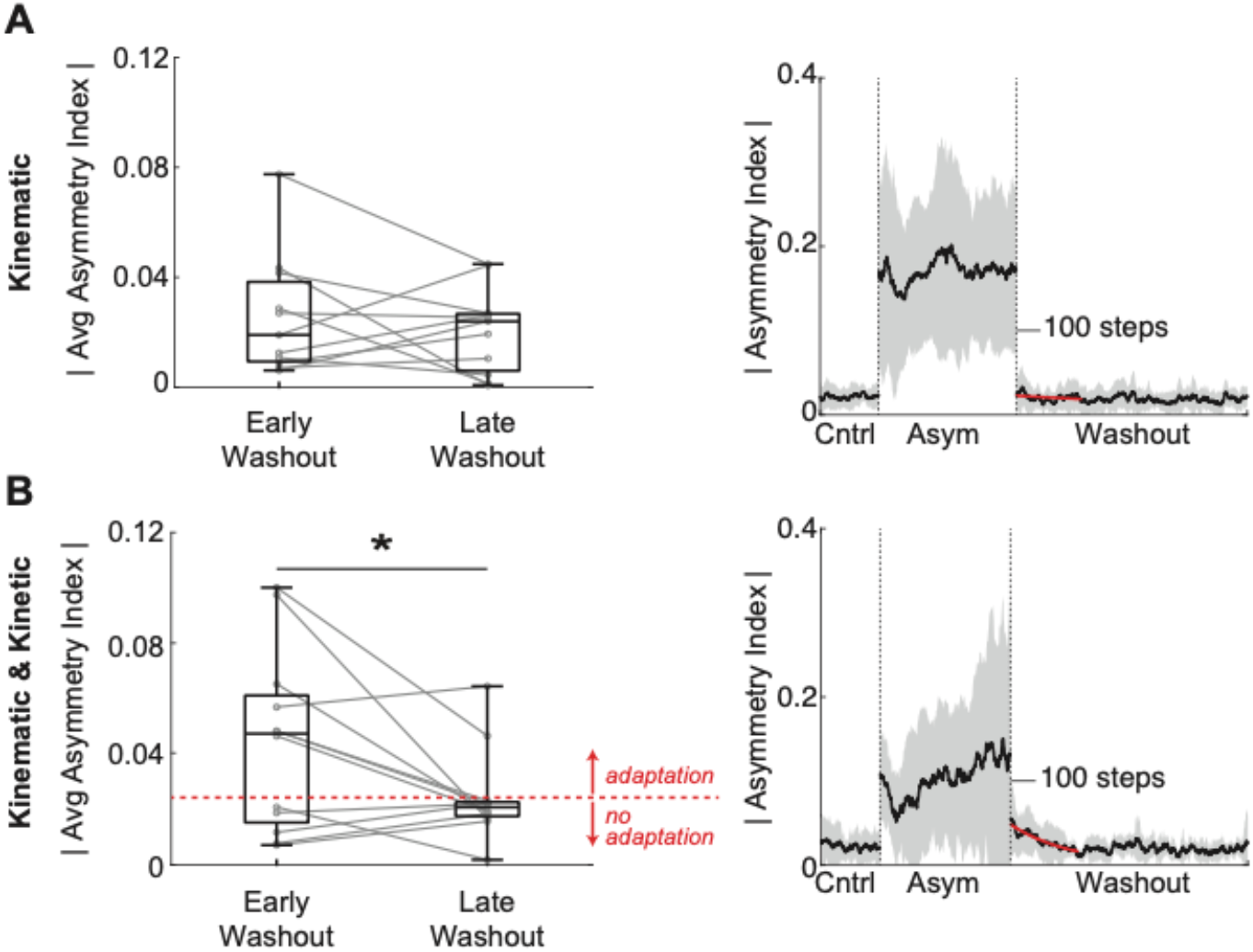
Results of the aftereffects analysis for Group 1 (kinematic) and Group 2 (kinematic and kinetic). (Left) Comparison of the absolute value of the average asymmetric index across participants in early (first 20 steps) and late (last 20 steps) washout. Red dotted line indicates the mean of the B1 late washout condition, representing the threshold for whether an individual participant showed evidence of de-adaptation. Thin gray lines show data for individual participants.* indicates a significant difference with p *<* 0.05. (Right) Absolute value of the average asymmetric index (AI) calculated from double stance times across each condition. AI = 0 represents symmetric gait. Gray shaded areas indicate standard deviation across participants. Red lines highlight the decay in asymmetry related to adaptation

For Group 2, the data was also not normally distributed (early, p = 0.003, late, p = 0.003). However, unlike Group 1, the median for the early washout was significantly greater than the median for the late washout (0.047 and 0.021), passing a right-tailed Wilcoxon signed rank test (p = 0.032), see Figure 4B. Therefore, Group 2, as a whole, demonstrated de-adaptation. The average behavior of the individuals within Group 2 that showed adaptation (i.e., had average AI values in early washout greater than the average) supports this result. Like Group 1, participants were nearly symmetrical in the control condition with average AI values of 0.0227 ± 0.0038 and the imposed asymmetry is evident in the asymmetric condition with average AI values of 0.1028 ± 0.0225. However, in the washout period, the presence of aftereffects from the imposed asymmetry is evident, indicated by the exponential decay in asymmetry in the beginning of the washout condition before the return to near baseline average AI values of 0.0222 ± 0.0078.

## 5 DISCUSSION

In this study, we tested if the sensorimotor adaptation in locomotion has an error threshold that initiates a persistent adaptation. The existence of this element is supported by observations that all people may have small and persistent asymmetric stepping. Only large limb asymmetries, for example, walking on the split-belt treadmill with 1:2 or 1:3 belt speed differences induced persistent adaptations [16,17]. Thus, we expected to detect the presence of an asymmetry threshold at which human participants initiate motor adaptations. Our aim was to test whether motor adaptation mechanism required a threshold of nonlinearities in limb dynamics and behavioral task to trigger adaptation. This was tested by imposing a whole-limb stride limiting (kinematic) constraint using a passive orthosis or a combination of the kinematic and loading (kinetic) constraints that induced stepping asymmetry. We tested the following hypothesis: the increasing sensory interlimb asymmetry triggers a persistent locomotor asymmetry with aftereffects at high, but not low, error values. We found that a kinematic constraint alone was not sufficient and required an additional kinetic constraint to trigger persistent adaptations in most participants. The adaptation caused by limb stepping asymmetry and interlimb difference in loading resembled asymmetric limb preference.

Only the combination of kinematic and kinetic constraints developed a persistent adaptation with limb preference. This suggests that the control system has “on-demand” resilience coordinating a kinematic asymmetry without the need for persistent changes. The result of testing for the presence of adaption under different conditions is summarized in Figure 4. Similarly, previous studies showed that small interlimb speed differences during split-belt treadmill locomotion did not induce adaptation, while the large differences did lead to adaptation with aftereffects (Yokohama et al., 2018). The increase in limb speed is positively correlated with the increase in loading, which can even be reliably used to identify limb speed in self-paced applications [30]. This would also support the expectation that large interlimb speed differences result in a large difference in interlimb loading, thus, creating a kinetic imbalance.

Why could a combination of kinematic and kinetic asymmetries, not a kinematic asymmetry alone, be the key to initiating adaptation? Our results indicate the existence of a sensory threshold for the onset of adaptation even at small interlimb stepping asymmetries with the same interlimb speeds. When interlimb speed difference is small, it generates turning and walking along curved paths [24,25]. This behavior falls within the domain of typical heading direction control and should not initiate a motor adaptation, as it would be disruptive to the regular turning function. It is likely that further increase in the kinematic asymmetry may also be within the physiological repertoire of precise limb control, for example, in the context of stepping over obstacles. However, further significant deviations that include likely both kinetic and kinematic differences fall outside of the expected spatiotemporal set of typical movements. This increased deviation from the expected level of errors may trigger the onset of adaptation.

It is likely that both individual experience and morphometry may define subject-specific magnitude that determines when “on-demand” resilience to kinematic asymmetries would be insufficient. In our study, 58% of participants developed adaptation with the kinematic and kinetic constraints that promote limb preference. We expect that further increase in the magnitude of the asymmetry condition could have resulted in the adaptation observed in the non-responder group. In the locomotor system, which is known to be a state-dependent dynamical system with multiple regimes [31], there may exist multiple stable states with varying inter- and intra-limb coordination to accommodate variable external conditions. One extreme example is the pattern of rhythmogenic activity of isolated spinal cord, which could switch from customary extensor-dominated to atypical flexor-dominated pattern in response to changes in external drives [28]. Similarly, both cortical and spinal pathologies may lead to persistent locomotor adaptations or maladaptations that require further rehabilitation. Our finding of error threshold in adaptation is novel and consistent with the typical practice of exaggerating locomotor deficits to promote locomotor rehabilitation [32,33].

Limitations to our study pertained to the magnitude of certain parameters (e.g., walking speed, amount of step length constrained, and threshold for unloading) being consistent across participants despite individual differences in preferred walking speed, step length, and weight. Regarding walking speed, the speed of 1.0 m/s was used across all participants because some may have found a typical healthy preferred speed of 1.4 m/s to be too challenging during the imposed asymmetry period due to their individual morphometry and/or behavioral set. The threshold for triggering adaptation may be related to the perceived difficulty of this task; therefore, future work may normalize the walking speed during the task to the individual’s preferred speed. Similarly, within groups, the amount of step length constrained was consistent across participants due to the passive orthosis being the same distance away from the limb regardless of the individual’s typical step length. This may have also contributed to the perceived difficulty of the task and ultimately the threshold for triggering adaptation. Future work may normalize the configuration of the passive orthosis to the participant’s step length or height. Finally, the monitoring of the vertical ground reaction forces during the task to ensure proper unloading in the participants who experienced both kinematic and kinetic constraints was subjective to the experimenter. Future work may automate this process and normalize the threshold to be considered unloading to the participant’s weight.

In populations with gait asymmetries due to injuries to the central nervous system (e.g., stroke and spinal cord injury), we can use motor adaptation principles to gradually reduce sensory error and learn a new desired movement. Before we can fully implement these principles into rehabilitation techniques, we must first learn how to effectively engage the motor adaptation mechanism. In this study, we found that both kinematic and kinetic components are needed in a paradigm to engage the mechanism and induce adaptation, an aspect that was not previously explored.

## Notes

### Competing Interest Statement

The authors have declared no competing interest.

